# Subtype-specific enhancer RNAs define transcriptional regulators and prognosis in breast cancers

**DOI:** 10.1101/2025.02.18.638890

**Authors:** Aamena Y Patel, Peyman Zarrineh, Jigar H Sheth, Sumitra Mohan, Mudassar Iqbal, Sankari Nagarajan

**Author notes:** Co-first authors (Aamena Y Patel, Peyman Zarrineh).

## Abstract

Gene expression is tightly controlled by DNA elements called enhancers by associating with lineage-specific transcription factors. These enhancers transcribe non-coding RNAs (called enhancer RNAs or eRNAs). eRNA expression is an early indicator of transcription factor activity and is associated with treatment response and survival in cancer patients. However, the attempts to identify prognostic eRNAs in breast cancers were inadequate, as these studies ignored the heterogenous nature of breast cancers with distinct molecular subtypes. By analysing ∼300,000 eRNA loci profiled using RNA-sequencing datasets from 1,095 breast cancer patients using machine learning approaches, we categorised eRNAs which are specific to breast cancer molecular subtypes and survival. The classified eRNAs were associated with gene pathways related to relevant subtypes. Interestingly, transcription factor analyses highlighted involvement of nuclear receptors other than the estrogen receptor with luminal-specific eRNAs. Basal eRNAs showed association with the transcriptional corepressor TRIM28 and androgen receptor. Luminal eRNAs were associated with better outcomes and Her2 eRNAs with worse outcome in patients. Overall, we demonstrate that machine learning approaches performed on RNA-seq datasets can classify subtype-specific and prognostic eRNAs which can be used to identify critical gene pathways and transcription factor networks in breast cancer.

## Introduction

Breast cancer is a highly heterogenous disease representing various subtypes with differences in the expression of classical hormonal and growth factor receptors, anatomical origin and gene alterations. Based on the expression of receptors-estrogen receptor (ER), progesterone receptor (PR) and human epidermal growth factor receptor-2 (Her2), pathologists classify breast tumours into different subtypes to enable targeted therapies. The molecular subtypes are hence classified as Luminal A (ER+ PR+ Her2-), Luminal B (ER+ PR+ Her2+), Her2 (Her2+ alone), basal (majorly triple negative ER-PR-Her2-) and normal-like [1,2]. The prognosis of each subtype-specific patients is distinct [3], emphasising the importance of these stratifications. LumA patients show a better prognosis than LumB while Her2 and basal patients have the worst outcomes. Overall, this highlights the importance of stratification of breast cancer patients at the molecular level to obtain efficient therapeutic benefits and better survival. Furthermore, breast tumours originate from either ducts and/or milk lobules in the breast. Lobular cancers are least studied breast cancers and have worse prognosis than ductal cancers. Despite the histological changes in these cancers, identifying their molecular differences is cumbersome, hindering the development of specific therapeutic strategies.

Gene transcription is highly controlled by *cis*-regulatory elements called enhancers. These are regions which can be located several 100s-1000s of kilobases away from gene promoters but can promote gene expression via 3D chromosomal interactions. Enhancers are bound and driven by lineage-specific transcription factors. Various studies in breast cancers showed expression of non-coding RNAs from highly active enhancers, termed as enhancer RNAs (eRNAs) [4–6]. eRNA expression is highly correlated with the activity of transcription factors such as ERα [4,5], FOXA1 [7], AR [8] and p53 [9]. Hence, enhancers which produce eRNAs can be evaluated to identify the transcription factors which are bound to and involved in the activity of those regions in an unbiased manner [10]. However, identifying eRNAs in patient samples is difficult, as the majority of eRNAs are non-polyadenylated and therefore are unstable. Hence, their detection requires high sample input and laborious techniques. As such, developing robust assays to identify eRNA expression in patient samples would be highly beneficial in developing patient-specific molecular signatures and key transcription factor networks.

Existing pan-cancer studies based on deeply sequenced and aggregated RNA-seq datasets identified cancer type-specific polyadenylated eRNAs which are more prognostic than mRNAs in certain types of cancer [11,12]. These eRNAs represented strong super-enhancer activity and can determine immunotherapeutic response in a cell-specific manner, resolving intra-tumour heterogeneity. Altogether, these findings suggests that eRNAs identified from RNA-seq datasets can provide sufficient power to be associated with patient survival. Despite the identification of 326 prognostic eRNAs in breast cancer from these studies, they are less prognostic than mRNAs. However, the heterogenous nature of breast cancers had not been considered in their analyses and this warrants further investigation following appropriate classification of patient datasets based on molecular subtypes.

In this study, we employed machine learning approaches on 302,951 eRNA loci identified from RNA-seq datasets from 1,095 breast cancer patient samples from previous studies [11,12], to refine eRNAs specific to molecular subtypes and survival in breast cancers. In addition to the major differences observed in molecular subtypes, our work identified that eRNAs are associated with key gene pathways and transcription factors specific to each subtype, highlighting the utility of stratified approaches on eRNA expression for understanding the important pathways involved in cancer progression.

## Materials and Methods

### Datasets

eRNA expression datasets from 1,095 breast cancer patient samples were downloaded from TCGA eRNA atlas (TCeA) platform [12]. These were mapped on 302,951 enhancer loci identified from H3K27ac chromatin immunoprecipitation (ChIP)-sequencing datasets from Hnisz *et al*., [13]. The patient metadata was downloaded from TCGA website https://portal.gdc.cancer.gov/projects/TCGA-BRCA. 90 and 49 samples expressing extremely low and high levels of eRNA respectively were filtered using outlier detection method with median-absolute-deviation (MAD) using *Scater* package (v1.22.0) [14], with number of MADs away from median required (nmads) as 1.5. Data from 975 breast cancer tumour samples were used to detect eRNAs which can distinguish between subtypes or survival.

For the Cistrome-based binding overlap analysis [15], data was downloaded from the section “TF ChIP-seq signals on eRNA loci” under “Integrated analysis” from TCeA. ChIP-seq and GRO-seq datasets were downloaded from published datasets for ER [16,17], H3K27ac [18], RNA polymerase [19] and GRO-seq [4].

### Subtype-specific eRNA detection, dimension reduction, and classification

Binary values (expressed or non-expressed eRNAs) were generated by applying *k*-means binning with k=2 on log2 rpkm eRNA expression values. *FSelector* (v0.34) [20] package was used to calculate the information gain values associated with the features (eRNAs) across the comparison sets (subtypes) and 0.05 cutoff was used for the binary expression values. As Her2 survival specific eRNAs with 0.05 cutoff were over 7000 loci, stringent cutoff (0.1) was used for survival-specific classification (living vs deceased patients as good vs poor outcome based on overall survival status of the patients). Average log2-transformed mean-centred rpkm values of eRNA (log mean centring or Logmc) over 1.0 cutoff were selected as continuous values.

*umap* (v.0.2.10.0) [21] was used to visualize UMAP plots. *randomForest* (v4.7-1.1) [22,23] was used to perform classifications into subtypes or survival status. The datasets were divided into 70% training and 30% test sets.

The random forest classifications were statistically assessed using different sets of subtype-or survival-specific eRNAs. These statistics described below use true positive (TP), true negative (TN), false positive (FP), and false negative (FN) metrics:

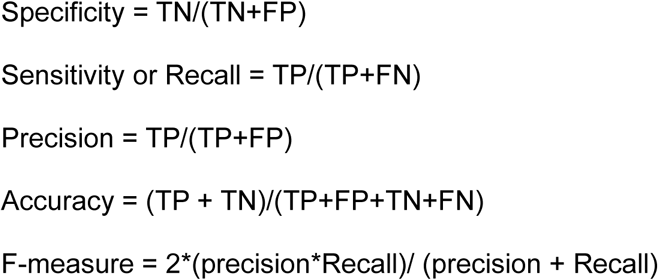

### Transcription factor and pathway analysis

To find the factors which significantly overlap with the eRNA loci, we overlapped the classified eRNAs with the eRNA loci annotated with transcription factor binding from Cistrome-based ChIP-seq datasets. Total unique binding sites were calculated for each factor irrespective of data from different cell lines. p-values were calculated using *phyper* function from *Hypergeometric distribution* in R stats (v4.4.2).

For motif enrichment analysis, *AME* (Analysis of Motif Enrichment) v5.5.7 [24] was used. Bed files were created with ±100bp flanks from the subtype-specific eRNA summits. These were compared to motifs in the *JASPAR 2022* Core vertebrates (non-redundant) database. For the background, the default shuffled input sequences were used.

*GREAT* (v3.0.0) [25] analysis was performed to obtain enriched pathways (perturbations) from *Molecular signature database* (MSigDB) [26] and *GO: Cellular Component* [27,28] associated with the genes close to the classified eRNA regions (any gene physically located up to 1 MB close to the eRNA loci). Hypergeometric FDR q-values and genome fraction values were plotted.

### KAS-sequencing and analyses

Low input N3-kethoxal-assisted single stranded DNA sequencing (KAS-seq) protocol was performed as described in Lyu *et al*., 2022 [29]. MCF7 cells (n=2) were cultured with DMEM with 10% fetal bovine serum, penicillin, streptomycin and 2 mM glutamine in a 12-well plate and labelled with 5 mM N3-kethoxal for 10 minutes. After labelling, 10,000 labelled cells were collected for further reactions. For tagmentation reaction, 50 ng of biotinylated DNA and 7.5 µl of the enzyme from Illumina Tagment DNA TDE1 enzyme and buffer kit were used. The transpositioned and amplified DNA samples were sequenced using NovaSeq 6000 with 1% PhiX spike-in generating paired-end reads. The reads were mapped on hg38 using bowtie2 [30] and *MACS2* (v2.2.7.1) broadpeaks option was used for peak calling [31]. *deeptools* (v3.5.1) [32] was used to construct bigwig files with rpkm settings (*bamCoverage*), average density plots (*computeMatrix* and *plotProfile*) and heatmaps (*computeMatrix* and *plotHeatmap*). Distal KAS-seq positive regions were calculated from all KAS-seq peaks ±5kb away from gene promoters and gene bodies.

### Survival analysis

Kaplan-Meier plots were performed using *survival* (v3.7-0) [33,34] and visualised with *survminer* (v0.4.9) [35] and *ggplot2* (v3.5.1) [36] in R (v4.4.1). p-values were calculated using log-rank test. Expression levels of subtype/survival-specific eRNAs were categorised as high or low by separating average of eRNA expression among patients as above or below 0 respectively. For basal subtype, highly expressed eRNAs were considered and thus, median of the average eRNA expression was used to stratify patients into high and low expression groups.

### Visualisation

*tidyverse* (v2.0.0) [37] was utilised for data manipulation in R (v4.4.2). Heatmaps were made using *ComplexHeatmap* (v2.22.0) in R with heatmap annotations [38,39]. *ggplot2* (v3.5.1) [36] and *cowplot* (v1.1.3) [40] was used to generate dotplots. Hierarchical clustering was performed using *hclust* function from R stats package (v4.4.2). The dendrograms were visualised using *ggtree* (v3.14.0) [41] and joined using *patchwork* (v1.3.0) [42]. Venn diagrams were created using *eulerr* (v7.0.2) [43,44].

## Results

### Identification of subtype-specific eRNAs

Previous work by Chen *et al*., [11] identified weak association of eRNA expression to patient survival in breast cancers. Given the heterogeneity and different subtypes of breast cancer, we hypothesised that eRNAs can associate with patient survival depending on subtype. Hence, we attempted to define eRNAs based on molecular subtypes (basal, luminal A/B and Her2) and histological tissue type (ductal vs lobular) and relate these subtype-specific eRNAs to survival. We reanalysed the datasets from Chen *et al*., [12] in which they mapped the eRNA signals in reads per kilobase per million mapped reads(rpkm) from 1,095 breast tumour samples on super enhancer loci (n=∼300K loci) based on H3K27ac occupancy datasets [13]. We generated two different measurements from the rpkm values of eRNA expression: (1) log-mean centring (Logmc) and (2) information gain (InfoGain). Based on given eRNAs from both measures, we performed subtype specific classification using random forest to assess the efficiency of eRNAs in distinguishing between subtypes (Fig. 1A). We observed that the overall metrics evaluating the performance of the classification were slightly better for InfoGain-identified eRNAs (Fig. 1B). However, sensitivity and F-measure (which measures predictive performance) were poorer for Her2 eRNAs, partly due to low number of Her2+ patients. The InfoGain-based approach identified larger set of eRNAs, compared to that of Logmc (Fig. 1C) and 75% of Logmc eRNAs overlap with InfoGain eRNAs (Fig. S1A). Basal eRNAs were identified more than luminal eRNAs using both measures and LumB-specific eRNAs could only be identified with Logmc (Fig. 1D). Uniform Manifold Approximation and Projection for Dimension Reduction (UMAP) visualisation of the data based on the classified eRNAs showed a clear separation of basal and luminal subtypes using both measures without distinguishing LumA/B (Fig. 1E). Both measurement-derived Her2 eRNAs clustered their patients closer to luminal patients, but in between luminal and basal patients. Surprisingly, neither measures could classify any distinct eRNAs for invasive ductal vs lobular cancer samples (Fig. S1B).

**Figure 1:**
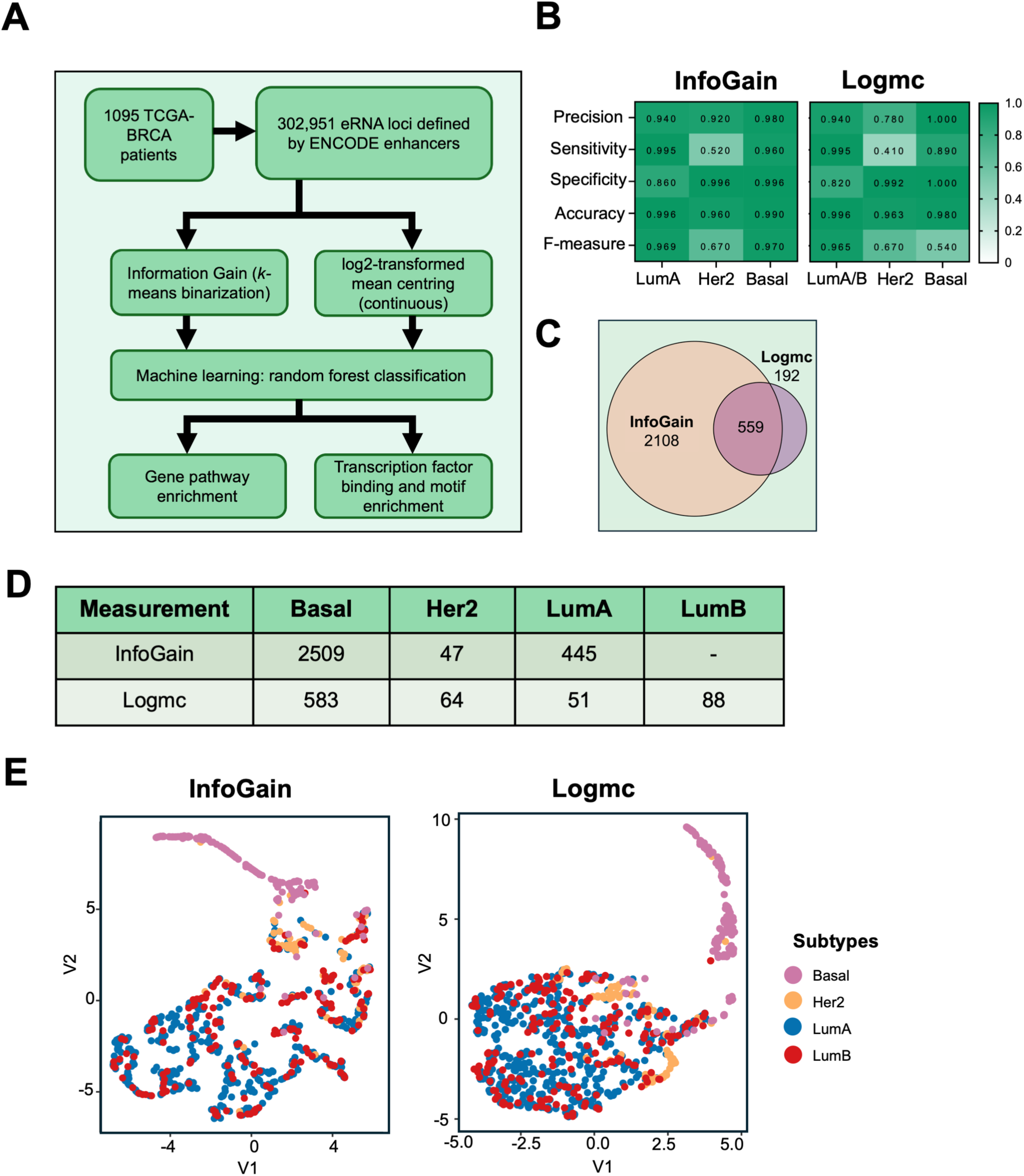
**(A)** Details of the analyses performed in this study. TCGA-BRCA – Breast Cancer RNA-seq datasets from The Cancer Genome Atlas (n=1,095). 302,951 enhancer regions defined by ENCODE H3K27ac datasets were utilised. *k*-means binning was used to convert the rpkm values to binarized values and information gain values were calculated. Log2-transformed centring of values to average expression (Logmc) provided continuous values. **(B)** Heatmap showing the statistics measures such as precision, sensitivity, specificity, accuracy and F-measure for information gain (InfoGain) and log2-transformed mean centring (Logmc) measurements classifying each subtype (luminal A/B, Her2 and basal). **(C)** Venn diagram representing the overlap of all InfoGain and Logmc-derived eRNA regions. **(D)** Number of eRNA regions classified per subtype with each measurement is shown in a table. Luminal B-specific regions could not be identified with InfoGain measure. **(E)** UMAP analysis showing the efficiency of InfoGain-(top 2 Principal Components) and Logmc-(top 4 Principal Components) derived eRNAs in classifying the clusters of patients from each subtype.

To visualise and verify the efficiency of the classification approach, we generated heatmaps with hierarchical clustering (Fig. 2A, S2A). Majority of the basal subtype patients were clustered together based on InfoGain-derived eRNAs, but Logmc-derived eRNA based clustering produced two groups of basal patients. Highly expressed eRNAs in patients with basal subtype had low expression in luminal subtype and vice versa. InfoGain measure could clearly classify both high/low expressed basal eRNAs. However, Logmc-derived basal eRNAs were highly expressed in most patients, but with mixed levels of expression in approximately 25% of patients (Fig. 2B-C). Hence, we could classify the InfoGain-derived eRNAs in basal subtype as high and low. While we observed a weak separation of high and low expressed InfoGain-defined LumA eRNAs, we could not distinguish their expression levels as strong as that of basal-specific eRNAs (Fig. S2B-C). Her2 specific eRNAs were highly expressed in approximately two-thirds of patients (Fig. S2D-E).

**Figure 2:**
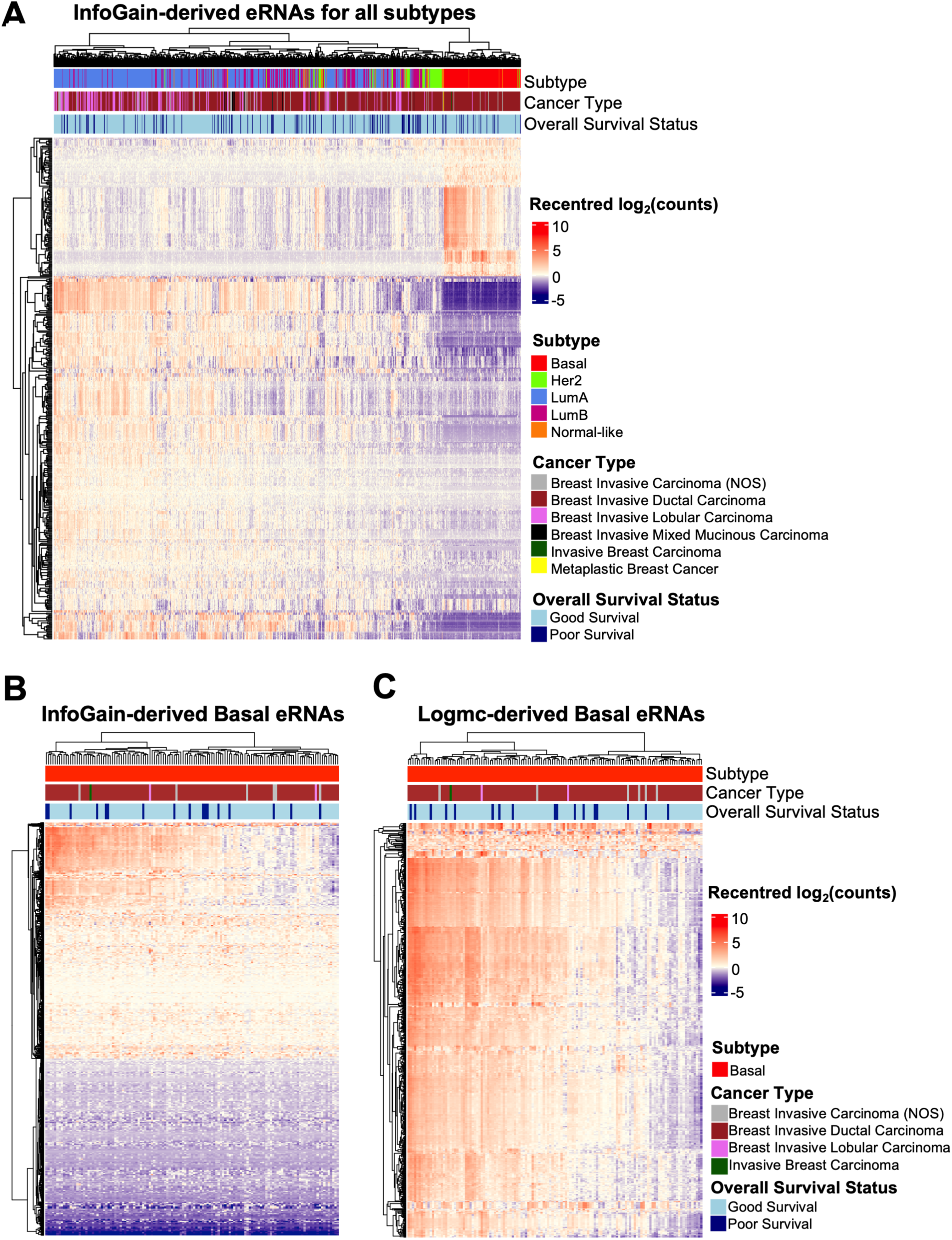
**(A)** Heatmap showing eRNA expression in log2-transformed mean-recentred values of breast cancer patient samples from TCGA on InfoGain-derived eRNA regions from all subtypes. Annotation of each patient with molecular subtypes, histological origin-based cancer type (cancer type) and overall survival status is shown. **(B-C)** Heatmap showing eRNA expression in log2-transformed mean-recentred values of basal breast cancer patient samples for InfoGain **(B)** and Logmc **(C)**-derived eRNA regions of basal subtype. Annotation of each patient with molecular subtypes, cancer type and overall survival status is shown.

Altogether, both measurements classify eRNAs efficiently based on subtypes, InfoGain allowed us to further distinguish samples based on high and low expression of eRNAs for basal subtype and performed better in statistical metrics. Hence, we focused on InfoGain-defined eRNAs for further analyses.

### eRNAs are associated with subtype-specific gene pathways

In order to verify the biological importance of the classified eRNAs per subtype, we examined the genes close to InfoGain-defined eRNA regions for pathway enrichment using GREAT analysis [25] with annotated pathways from Molecular Signature database [26]. We observed that basal high eRNAs were associated with basal-specific pathways or luminal/ER downregulated pathways and vice versa (Fig. 3A). Immunological and development associated signatures such as T-cell/FOXP3 and puberty-related pathways were enriched with basal high eRNAs. Bone relapse-specific downregulated genes and SMAD2/SMAD3 and MYB target genes were also associated with basal high eRNAs, while brain relapse and MYC (ER cofactor and ER target gene)-specific downregulated genes were associated with basal low eRNAs. Endocrine-resistant pathways which denote the luminal ER response markers were enriched with basal low eRNAs. Luminal eRNAs identified ER/luminal upregulated or basal downregulated pathways. Consistently, brain relapse downregulated and bone relapse upregulated pathways are associated with luminal eRNAs.

**Figure 3:**
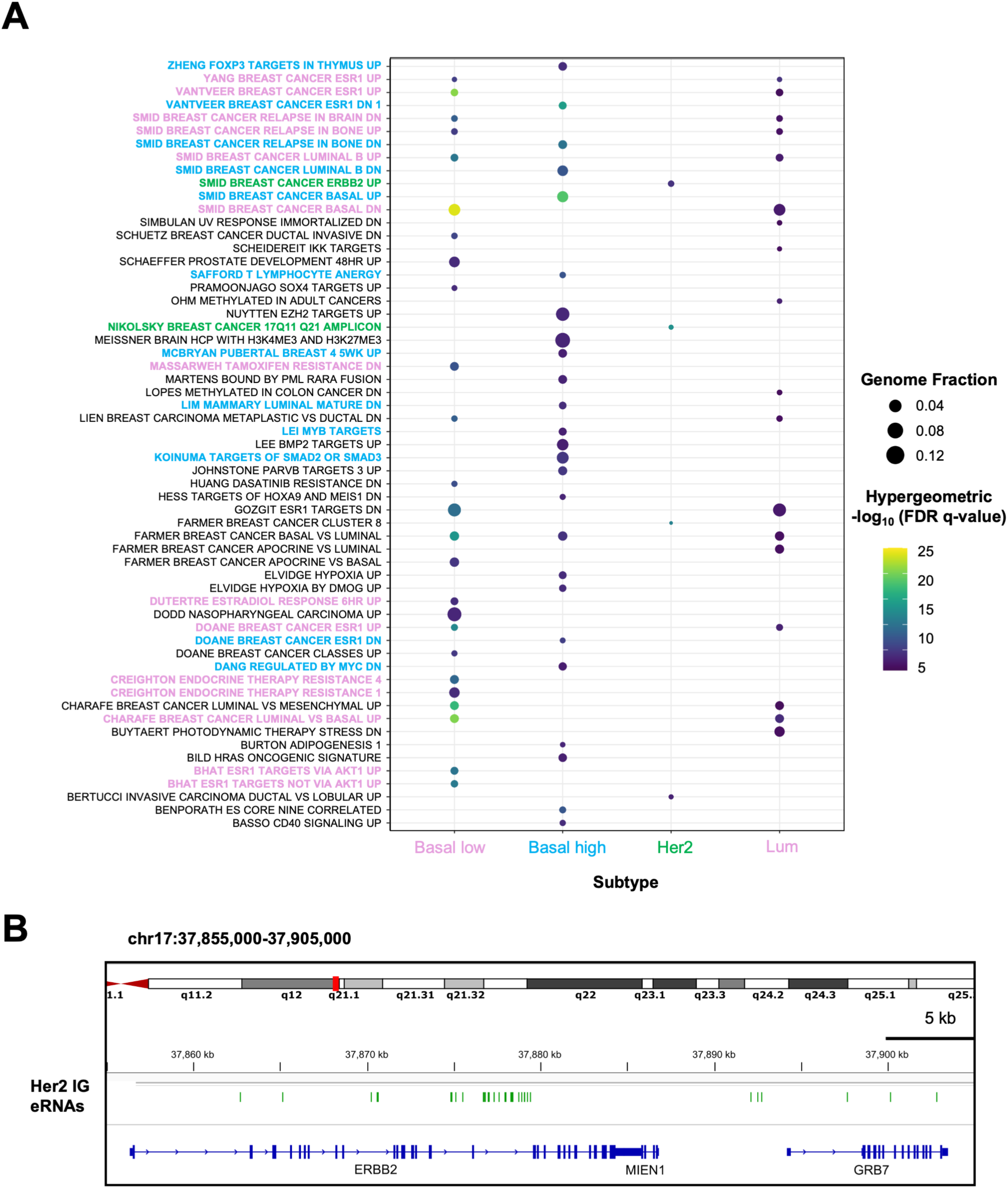
**(A-C)** Dotplots showing gene pathway analyses performed on InfoGain-derived eRNA regions of low expression (Basal low) and highly expression (Basal high) in basal subtype, luminal A and Her2-specific regions using GREAT tool and MSigDB database. Colour keys represent hypergeometric-log10 FDR q-value and genome fraction (ratio of observed genes by total number of genes in a pathway gene set). Pathway names were highlighted in red (luminal or basal low), blue (basal high) and green (Her2) specific to subtypes. Only significant pathways are shown. Pathways specific to luminal, ER-regulated, basal, Her2, endocrine resistance, relapse, immune and development-related signatures are highlighted. **(B)** Multiple gene view around q11-21 amplified region from chromosome 17 close to *ERBB2* (Her2) gene with the Her2-specific eRNA regions derived from InfoGain (green).

Many ER target/luminal genes (*ESR1*, *MLPH*, *GATA3*, *CT62/THSD4, XBP1*) enriched in these pathways are lineage-specific factors reportedly upregulated in ER+ breast cancers [45–47]. To verify if the eRNA loci close to these genes are actively bound and regulated by ER in patients, we examined ER binding using published ER ChIP-sequencing datasets from drug responsive (MCF7, ZR-75-1) and resistant (BT474, MCF7 TamR Tamoxifen resistant) ER+ cell lines, primary patient samples with differential survival status (good and poor outcome) and metastatic samples [17]. We identified that eRNA regions close to the above-mentioned genes also present with proximal ER binding in most of the patient samples and cell lines (Fig. S3).

Her2-specific eRNAs showed enrichment of pathways related to *ERBB2* (gene of Her2) (Fig. 3A). The pathway ID “NIKOLSKY_BREAST_CANCER_17Q11_Q21_AMPLICON” represents genes from chromosome region 17q11-21, and part of this region harbours the genes *ERBB2* and *MIEN1* which are activated/amplified in Her2-positive cancers (Fig. 3B). There were 30 Her2-specific eRNA regions in the vicinity of these genes. This suggests that our analysis identified relevant eRNA loci for Her2-type breast cancers, but caution should be employed to rule out any possible patient-specific copy number amplifications confounding the eRNA expression levels. Altogether, our pathway analyses suggest that eRNA expression can be associated directly with subtype/lineage-specific gene pathways.

### Subtype-specific eRNAs are associated with key transcription factors and epigenetic regulators

To identify the important factors associated with the classified eRNAs, we integrated them with the published ChIP-seq datasets of transcription factors and epigenetic regulators from Cistrome platform [15] (Fig. 4A). Basal regions looked distinct with overlap on Tripartite motif-containing transcriptional corepressor TRIM28, histone variant H2AZ which usually marks active enhancers and androgen receptor (AR). LumA subtype eRNA loci showed significant enrichment of known key players such as nuclear receptors glucocorticoid receptor (GR) [48] and peroxisome proliferator-activated receptor gamma (PPARG), and BRD4 (bromodomain and extra-terminal domain BET-containing epigenetic reader and active enhancer-associated protein [49]), etc. Surprisingly, ER was not significantly enriched in LumA regions, but forkhead domain containing protein FOXA1, which is a lineage specific factor of the LumA subtype is enriched only in basal low regions. Her2 subtype regions showed strong enrichment of several transcriptional regulators including general transcription factor IIi (GTF2I), CCAAT/enhancer-binding protein beta (CEBPB), Snail Family Transcriptional Repressor-2 (SNAI2), SWI/SNF Related BAF Chromatin Remodelling Complex Subunit ATPase-4 (SMARCA4, a major subunit of SWI/SNF chromatin remodelling complex) and zinc finger protein ZNF263. Our analyses also identified role of another nuclear receptor and dormancy marker NR2F1 in both basal and luminal sites.

**Figure 4:**
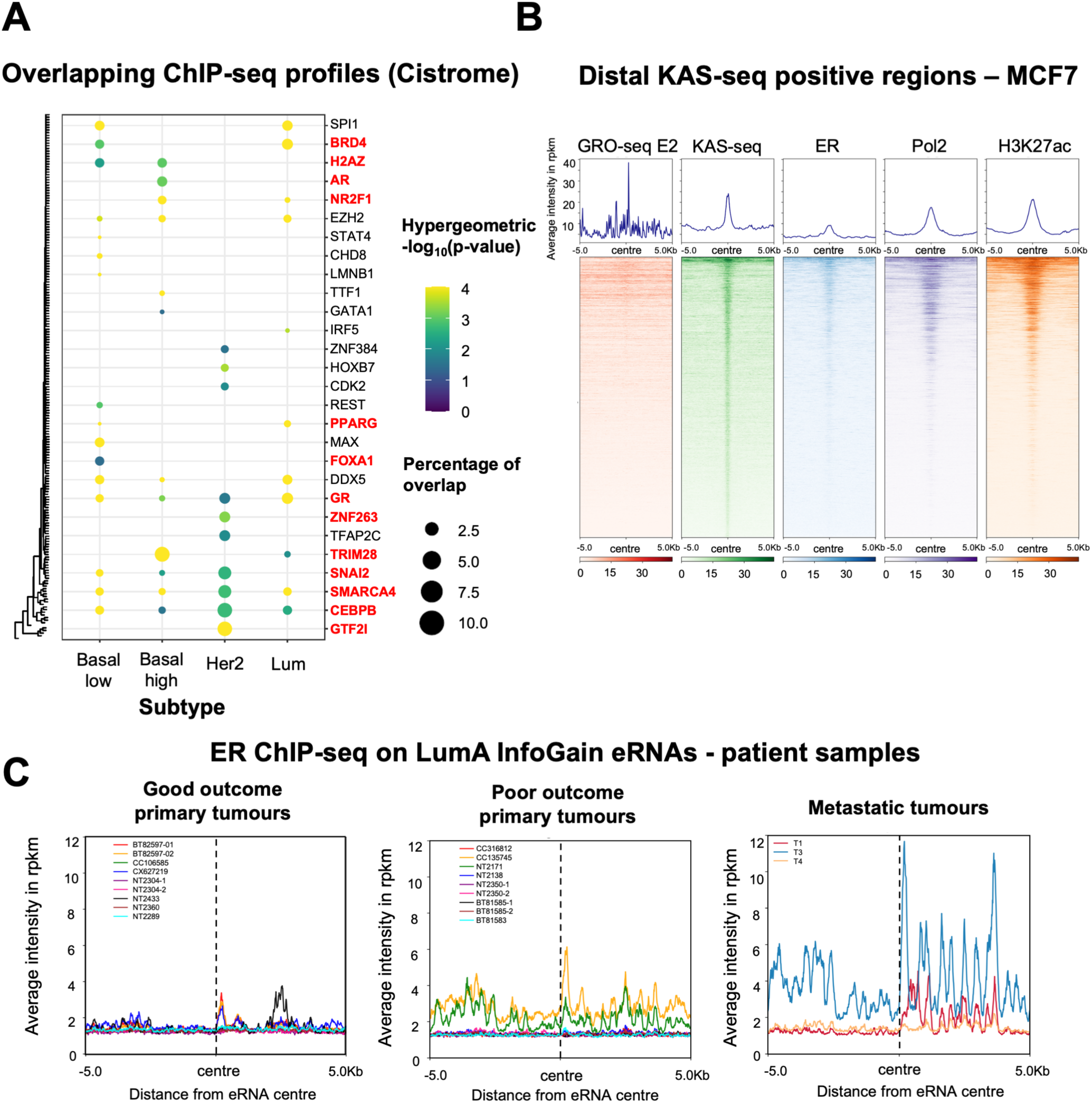
**(A)** Dotplot showing Cistrome-based overlap of binding with published ChIP-seq datasets performed on InfoGain-derived eRNA regions of low expression (Basal low) and highly expression (Basal high) in basal subtype, luminal A and Her2-specific regions. Colour keys represent hypergeometric-log10 p-value calculated by phyper function from hypergeometric distribution (hypergeometric package) from R and percentage of overlap of the eRNA regions with the published binding sites for each factor. Only significant factors are shown. Factors of interest are highlighted in red. **(B)** Average density plots and heatmaps showing the rpkm values of eRNA expression from global run-on sequencing (GRO-seq) from 40 minutes of estrogen treated MCF7 cells and kethoxal-assisted ssDNA (KAS)-sequencing, and ER, RNA polymerase II and H3K27ac occupancy from MCF7 cells which are continuously grown with serum with hormones. The regions are mapped ± 5 kb away from the centre of distal eRNA peaks (9,036 regions) defined by KAS-seq from MCF7 cells. **(C)** Average density plots showing the rpkm values of ER binding from patient samples with good outcome and poor outcome primary tumours and metastatic samples. The regions are mapped ± 5 kb away from the eRNA peak centre defined by luminal A-specific eRNAs classified by InfoGain. ER ChIP-seq datasets from patient samples were downloaded from Ross-Innes *et al*., 2012. Each line represents ER binding from different patients. Patient ids are provided in the legend. ER ChIP-seq data from patient ID 82590 was removed due to background.

Furthermore, we looked at the frequency of DNA motifs associated with transcription factors on the 100 bp flanked regions of the classified eRNAs (Fig. S4A). Consistent with the results from Cistrome analyses, we could not observe any direct motifs of ER. However, we identified binding motifs of key luminal factors such as cAMP response element binding protein (CREB) factors, forkhead domain family proteins, retinoic acid/retinoid X receptors RARA/RXR and PPARG. Binding motifs of AP-2 activator protein family, which play an important role as pioneer factors on ER-dependent luminal enhancers, were enriched on basal low regions. Binding motifs of regulators of differentiation (Myocyte enhancer factors MEF2A, MEF2B, MEF2C and MEF2D) were observed on eRNAs from all the subtypes. Interferon regulatory clusters (IRF1, IRF4, IRF5, IRF8, IRF9) binding motifs were more enriched on Her2 regions. Motifs for homeodomain containing factors such as HD-CUT and HOX were specifically enriched on basal and Her2 regions. Overall, Cistrome binding overlap and motif analyses on eRNA loci suggest key transcription factors and epigenetic modulators specific to each subtype, more specifically the association of nuclear receptors other than estrogen receptor and forkhead proteins with luminal or basal low eRNAs/enhancers.

To validate the ER-independent nature of LumA eRNAs, we performed enhancer RNA profiling using kethoxal-assisted ssDNA sequencing (KAS-seq [50]) in MCF7 cell line which is an ER+ LumA type breast cancer cell line. This assay identifies regions of single stranded DNA (ssDNA) representing active transcription. We profiled the KAS-seq signals on distal sites 5 kb away from genes (9,036 distal KAS-seq positive regions) and integrated them with global run-on-based sequencing (GRO-seq to detect nascent RNA [51]) and ER, RNA polymerase-II, and H3K27ac occupancy (Fig. 4B). We observed that ER binding occupies only the strongest ssDNA and GRO-seq-positive sites. Furthermore, in MCF7 cells, we observed that ER is bound 1 kb away from the centre of InfoGain-derived LumA eRNA loci (Fig. S4B, S3).

Additionally, we integrated luminal eRNAs with ER ChIP-seq datasets from patient samples and we observed similar ER binding which is not on the centre of the eRNA regions (Fig. 4C, S3). These eRNA regions showed slight proximal/distal ER binding in 4/9 of the good outcome patients but showed strong ER binding in 2/3 of metastatic and 2/9 of poor outcome patients. This is consistent with the lack of overlap with ER binding (Fig. 4A) in Cistrome analysis, as Cistrome-based ER ChIP-seq datasets were majorly obtained from MCF7 cells which represent good outcome patients with effective treatment response. Overall, our findings suggest that not all the actively transcribing distal eRNA loci possess ER binding, but if they do, they present with ER binding few kilobases away, more prominently in patients with aggressive and metastatic tumours.

### InfoGain-derived eRNAs can be prognostic in breast cancers

Next, we investigated the prognostic power of subtype-specific eRNAs. Kaplan Meier curve-based survival analysis of LumA-specific eRNAs showed that patients with high expression of these eRNAs have better survival (Fig. 5A). This is consistent with expression of ER and its target genes being positively associated with survival in luminal patients [52]. Expression of Her2 and basal subtype-specific eRNAs did not show any significant association with patient survival (Fig. 5B-C).

**Figure 5:**
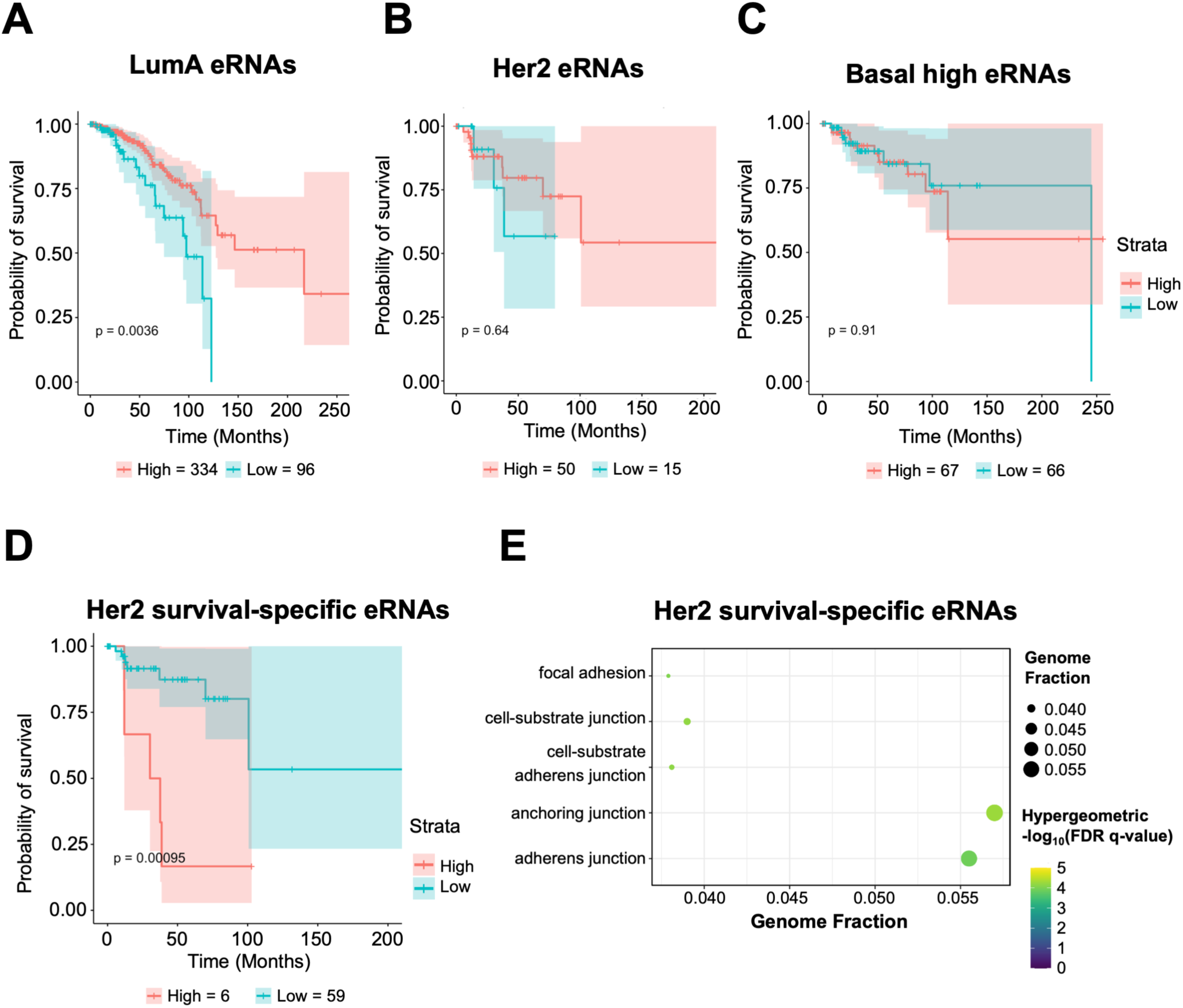
**(A-C)** Kaplan-Meier plots showing the probability of overall survival of patients with each subtype (luminal **(A)**, Her2 **(B)** and basal high **(C)** eRNAs) who show low and high average expression of InfoGain-derived subtype specific eRNAs among patients. Number of months of patient survival after diagnosis is shown. Plots represent 430 LumA patients, 65 Her2 patients and 133 Basal patients. p-values were calculated using log-rank test. Numbers of patients in each category (high and low average eRNA expression) are mentioned under each plot. **(D)** Kaplan-Meier plot showing the probability of overall survival of Her2-positive patients who show low and high average expression of InfoGain-derived subtype specific eRNAs among patients, classified based on survival (dead-poor survival or alive – good survival). Number of months of patient survival after diagnosis is shown. p-value was calculated using log-rank test. Numbers of patients in each category (high and low average eRNA expression) are mentioned under each plot. **(E)** Dotplot representing pathway enrichment analysis on Her2-specific InfoGain-derived eRNAs showing significant association with cell adhesion pathways. Colour keys represent hypergeometric-log10 FDR q-value and genome fraction (ratio of observed genes by total number of genes in a pathway gene set). Only significant pathways are shown.

To identify the subtype-specific prognostic eRNAs with high confidence, we performed random forest classification on both measures distinguishing patients with good or poor outcome per subtype. InfoGain measure successfully identified 342 eRNAs which classifies the survival status of Her2 subtype with 0.1 threshold (S5A-B), but not for other subtypes even with less stringent threshold. Survival and hierarchical clustering analyses showed that Her2-positive patients with high average expression of survival-specific eRNAs have worse outcomes (Fig. 5D, S5A). The genes close to these prognostic Her2 eRNAs are heavily enriched in cell adhesion/junction pathways (Fig. 5E), implying the importance of deregulation in extracellular matrix or epithelial-mesenchymal transition pathways in poor prognosis. Logmc measure could not identify any eRNAs specific to survival. Overall, our machine learning-based analyses employed on RNA-seq datasets could identify subtype and survival-specific eRNAs which are associated with differential outcome in breast cancer patients.

## Discussion

Our studies attempted to identify eRNAs which are specifically expressed in each subtype based on gene expression, anatomical origin and survival in breast cancer. We observed that eRNA expression in cancers of ductal and lobular origin as well as for LumA and LumB subtypes looks majorly similar, but our machine learning approach can identify subtype-specific eRNAs and luminal and Her2-specific prognostic eRNAs. Our integrated analyses identified eRNAs close to genes which are highly active in patient samples and key transcription factors and epigenetic proteins which can play important roles in lineage-specific breast cancers. Furthermore, our study identified nuclear receptors other than ER and pioneer factors which could be important in LumA breast cancers. Hence, our study supports the avenue of utilising eRNAs as lineage-specific biomarkers for identifying upstream regulators and prognosis.

Around 90% of eRNAs are bidirectional and non-polyadenylated [53]. TCGA expression datasets are based on RNA-seq assays, which capture only non-polyadenylated RNAs. Thus, analysing the expression of eRNAs on mRNA-seq datasets might not be adequate. However, non-polyadenylated transcripts are highly unstable leading to a low half-life of 2-3 mins for eRNAs [54]. Hence, detection of polyadenylated eRNAs on highly expressed enhancer regions can be easily adapted to any clinical lab for routine diagnostic purposes on FFPE samples where RNAs are heavily degraded. Further work on eRNA expression on patient samples using robust nascent RNA detection methods like ChRO-seq is still warranted to acquire strongest lineage-specific or prognostic eRNAs. However, this is a tedious process and currently there are no nascent RNA datasets available on breast cancer patient samples.

Our study identified similar subtype-specific eRNAs from different measures. To note, InfoGain-based analyses identified both high/low expressed and prognostic eRNAs. Hence, using InfoGain for machine learning-based classification of biomarkers would provide greater benefit. This approach can also be used to identify prognostic biomarkers in other cancers where their heterogenous nature is established.

We identified that lobular and ductal cancers are indistinct in eRNA expression, which is similar to previous studies with mRNA expression [55,56], even though there are many lobular cancer-enriched mutations in genes such as *CDH1*, *FOXA1* and *ARID1A*. Hence, understanding the drug resistance mechanisms to existing ER-targeted therapies like tamoxifen or developing lobular-specific drug targets are challenging. Various studies established that the tumour and immune microenvironment in lobular cancers are distinct in comparison to ductal cancers [57,58]. This emphasises the utility of single cell RNA-seq datasets performed on high stroma vs tumour content to be more appropriate for identifying specific molecular signatures in lobular cancers.

While ER motifs or overlap with ER binding could not be observed on exact loci of luminal eRNAs, these regions showed strong enrichment of other nuclear receptors like GR/RARA/RXRA/PPARG/NR2F1, forkhead or AP2 transcription factors. We identified eRNAs close to *ESR1* and other lineage-determining genes with strong distal ER binding, especially in patients with aggressive cancers. Altogether, this suggests that nuclear receptors other than ER coupled with the forkhead family and/or AP2 pioneer factors can play key roles on active enhancers in primary tumour samples. Furthermore, it is possible that ER can either associate with these factors distally in good outcome patients or it is reprogrammed to be more active in patients with metastasis or drug resistance. Interestingly, basal-specific eRNAs identified the involvement of AR, emphasising its role in luminal triple negative breast cancers [59], and the corepressor TRIM28 which promotes cancer stem cell proliferation and metastasis in triple negative breast cancers [60,61].

In the previous studies, Chen *et al*., [11,12] identified 326 prognostic eRNAs in breast cancers without subtyping them. Surprisingly, these eRNAs don’t overlap much with any eRNAs classified by our approaches and they show weak association with survival (Fig. S5C). Our InfoGain-based analyses showed that eRNA expression can be associated with good or poor prognosis dependent on subtype. This underscores the importance of molecular subtyping before identifying any prognostic biomarkers in heterogenous cancers. While eRNA expression is associated as a positive biomarker for therapeutic response to BET inhibitors and immune checkpoint inhibitors [11,62], high expression of genomic instability-specific eRNAs is related with worse survival in breast cancer patients [63]. Hence, identifying prognostic eRNAs based on subtypes using InfoGain measure or any other sophisticated machine learning approaches is critical. Further work on RNA-seq datasets with adequate number of patients across minor subtypes would be beneficial.

Altogether, our study emphasises the importance of subtyping to identify prognostic eRNAs/enhancers, molecular signatures and upstream regulators which can be therapeutically relevant and targetable. Our findings further highlight the possibility of constructing gene regulatory transcription factor networks related to patient survival by looking into classical RNA-seq datasets, as other laborious epigenetic analyses require patient samples in fresh frozen conditions which are extremely limited.

## Data and code availability

KAS-seq datasets are available upon request.

## Acknowledgements

We thank Danielle Olivier for the initial exploration of the datasets. We thank Prof. Andrew Sharrocks and Dr. Hayden Jones, University of Manchester and Claire Turner, Lobular Cancer UK charity for the critical reading and evaluation of the manuscript. SN, AP and SM are supported by CRUK Career Development award (RCCFEL\100069).

## Contributions

SN – conceptualization; AP, PZ, JHS, SN – methodology, software, validation, formal analysis, investigation, visualization; SM, MI – methodology; SN, MI – supervision; SN – writing original draft; AP, PZ, JHS, SM, MI, SN - review & editing of the manuscript; SN-funding acquisition.

## Supplementary figure legends

**Figure S1: (A)** Venn diagram showing the overlap of all InfoGain and Logmc-derived eRNA regions from each subtype. Luminal A eRNAs are used for the overlap, as InfoGain could not identify any luminal B specific eRNAs. **(B)** UMAP analysis showing the 4 top Principal Components (PC) in classifying the clusters of patients from each histological subtype-invasive ductal, invasive lobular and others which includes mixed histology (NOS), mucinous, medullary and metaplastic carcinomas. As the classification did not yield enough ductal or lobular-specific eRNAs (n=1), PC components were made based on all the eRNAs filtered out of outliers, to show their indistinguishable nature.

**Figure S2: (A)** Heatmap showing the eRNA expression in log2-transformed mean-recentred values of all breast cancer patient samples from TCGA on Logmc-derived eRNA regions from all subtypes. Annotation of each patient with molecular subtypes, histological origin-based cancer type (cancer type) and overall survival status is shown. **(B-E)** Heatmap showing eRNA expression in log2-transformed mean-recentred values of luminal A (**B, C**) and Her2 (**D, E**) patient samples, on InfoGain **(B, D)** and Logmc **(C, E)**-derived eRNA regions derived specific to the respective subtypes. Annotation of each patient with molecular subtypes, histological cancer type and overall survival status is shown.

**Figure S3:** Multiple gene view close to *ESR1* (ER) **(A)**, GATA3 **(B)**, MLPH **(C)** and THSD4/*CT62* **(D)** genes showing the occupancy of ER (blue rectangles) in good outcome and poor outcome primary tumour and metastatic samples from ER+ patients and cell lines representing drug sensitive (MCF7, ZR-75-1) and tamoxifen-resistant (BT474, MCF7 Tamoxifen resistant TamR) cell lines, on the eRNA regions derived from Logmc and InfoGain specific to luminal A subtype. ER ChIP-seq datasets from patient samples and cell lines were downloaded from Ross-Innes *et al*., 2012.

**Figure S4: (A)** Dotplot showing motif enrichment analyses performed on InfoGain-derived eRNA regions of low expression (Basal low) and highly expression (Basal high) in basal subtype, luminal A and Her2-specific regions. Colour keys represent log10 average of the ratio of true positives over false positive motifs and-log10 adjusted p-value from AME tool. Only significant pathways are shown. Transcription factor motifs of interest are highlighted in red. **(B)** Average density plot showing the rpkm values of ER occupancy from MCF7 cells grown in serum containing medium representing continued supply of hormones. The regions are mapped ± 5 kb away from the eRNA peak centre defined by luminal A-specific eRNAs derived from InfoGain. High – high ER binding regions, low – low/no ER binding regions.

**Figure S5: (A)** Heatmap showing eRNA expression in log2-transformed mean-recentred values on InfoGain-derived Her2 and survival-specific eRNA regions. Annotation of each patient with molecular subtypes, cancer type and overall survival status is shown. eRNA loci associated with good and poor survival are marked with arrows. (**B**) Table showing the statistics measures such as precision, sensitivity, specificity, accuracy and F-measure for information gain (InfoGain) measurements classifying Her2 subtype based on survival (good and poor outcome patients). **(C)** Heatmap showing eRNA expression in log2-transformed mean-recentred values in rpkm on prognostic eRNA loci (n= 326) identified by Chen *et al*., 2018. Annotation of each patient with molecular subtypes, cancer type and overall survival status is shown.

