## Supplementary figures for "Subtype-specific enhancer RNAs define transcriptional regulators and prognosis in breast cancers"

#### Figure S1

**A**

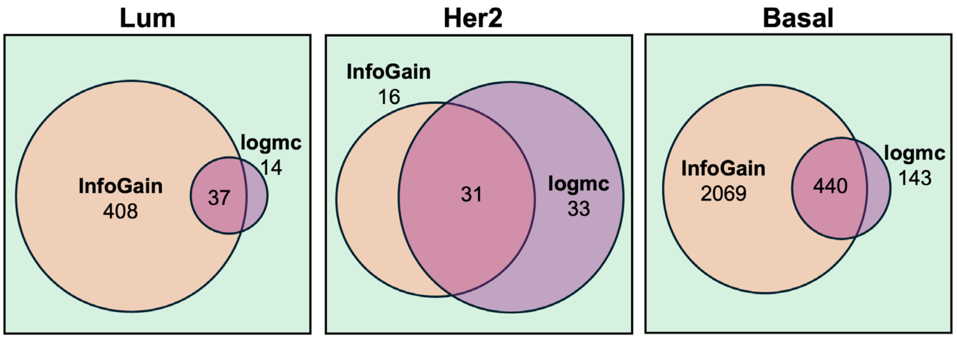


**B**

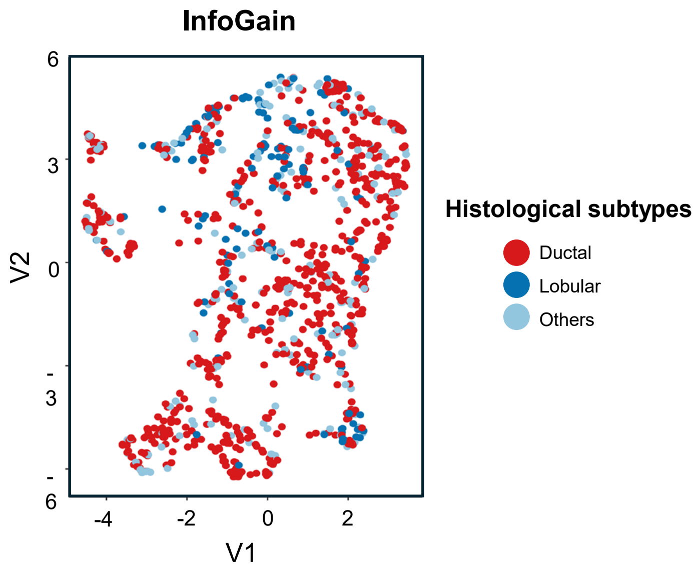

### Figure S2

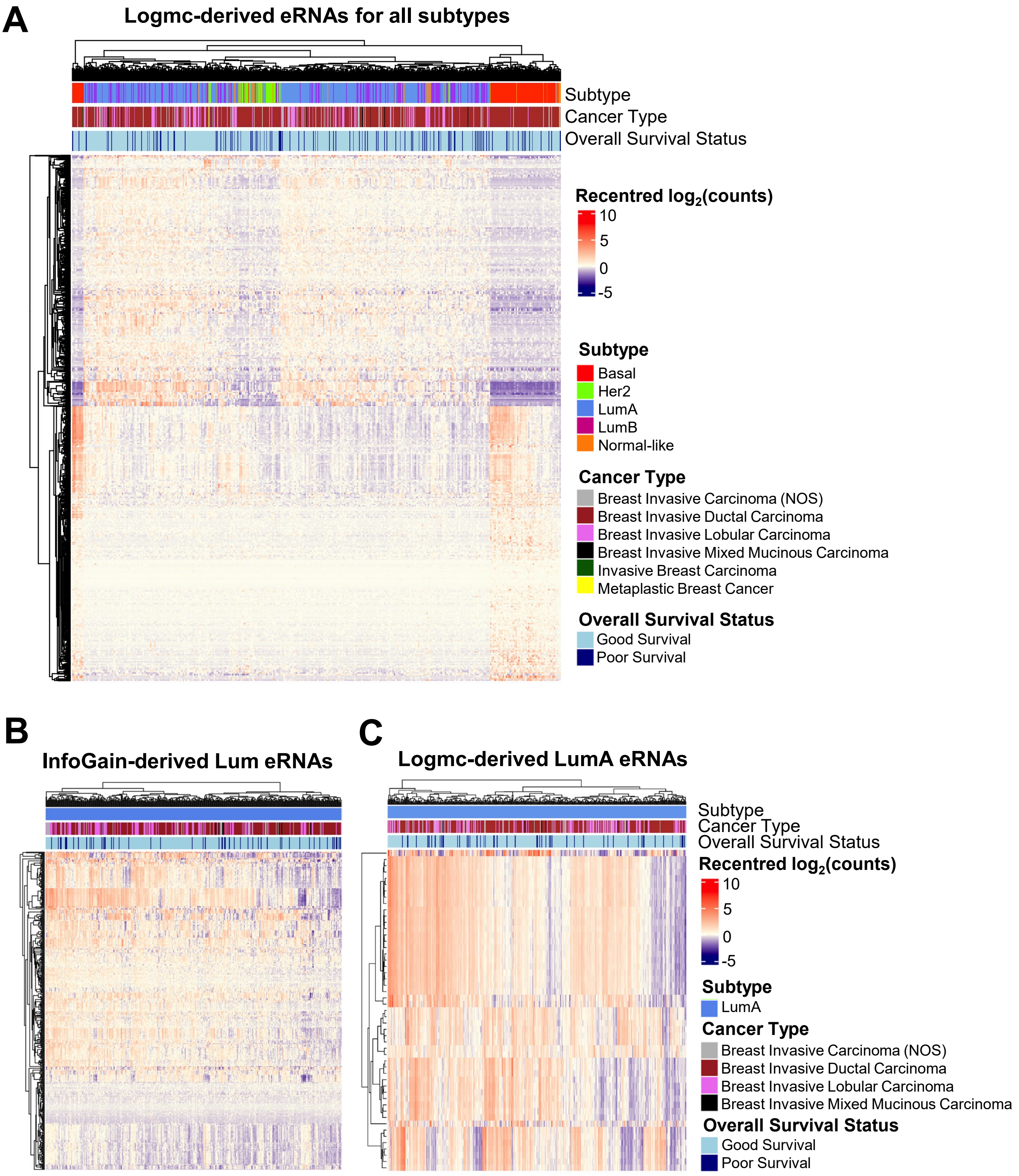
A
Logmc-derived eRNAs for all subtypes
Subtype
Cancer Type
Overall Survival Status
Recentred log2(counts)
10
5
0
-5
Subtype
Basal
Her2
LumA
LumB
Normal-like
Cancer Type
Breast Invasive Carcinoma (NOS)
Breast Invasive Ductal Carcinoma
Breast Invasive Lobular Carcinoma
Breast Invasive Mixed Mucinous Carcinoma
Invasive Breast Carcinoma
Metaplastic Breast Cancer
Overall Survival Status
Good Survival
Poor Survival
B
InfoGain-derived Lum eRNAs
C
Logmc-derived LumA eRNAs
Subtype
Cancer Type
Overall Survival Status
Recentred log2(counts)
10
5
0
-5
Subtype
LumA
Cancer Type
Breast Invasive Carcinoma (NOS)
Breast Invasive Ductal Carcinoma
Breast Invasive Lobular Carcinoma
Breast Invasive Mixed Mucinous Carcinoma
Overall Survival Status
Good Survival
Poor Survival

### Figure S2 cont.

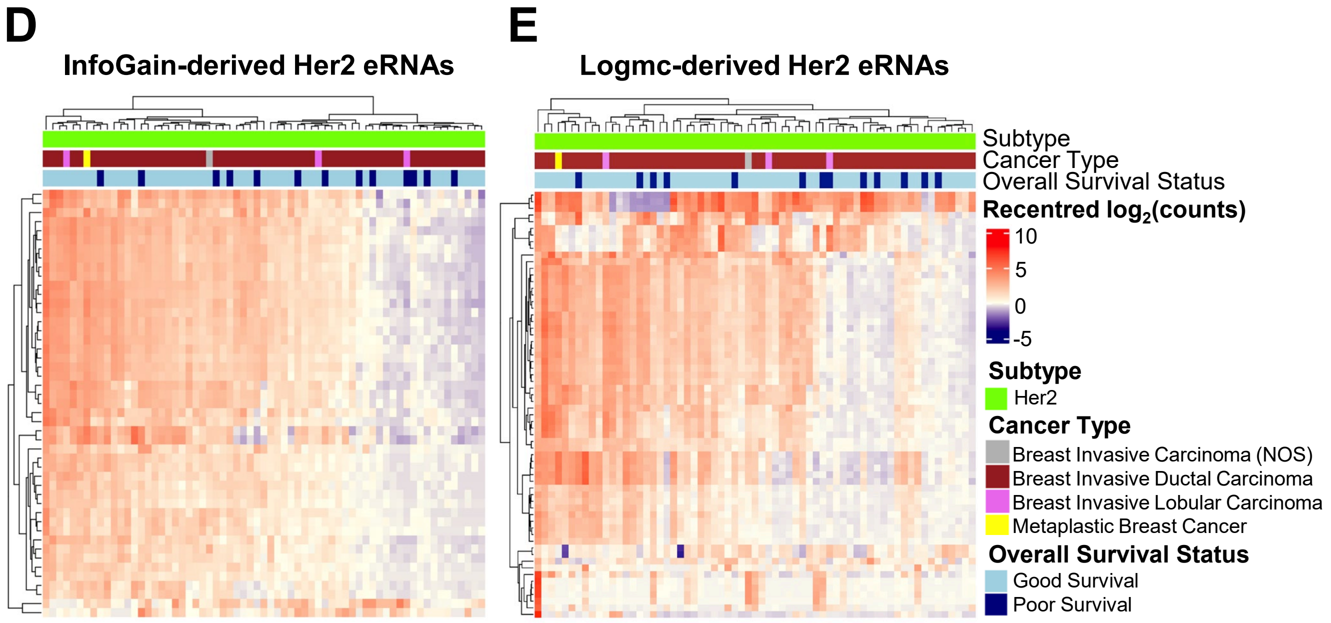

### Figure S3

**A**

**ER ChIP-seq from patient samples and cell lines on eRNA regions**

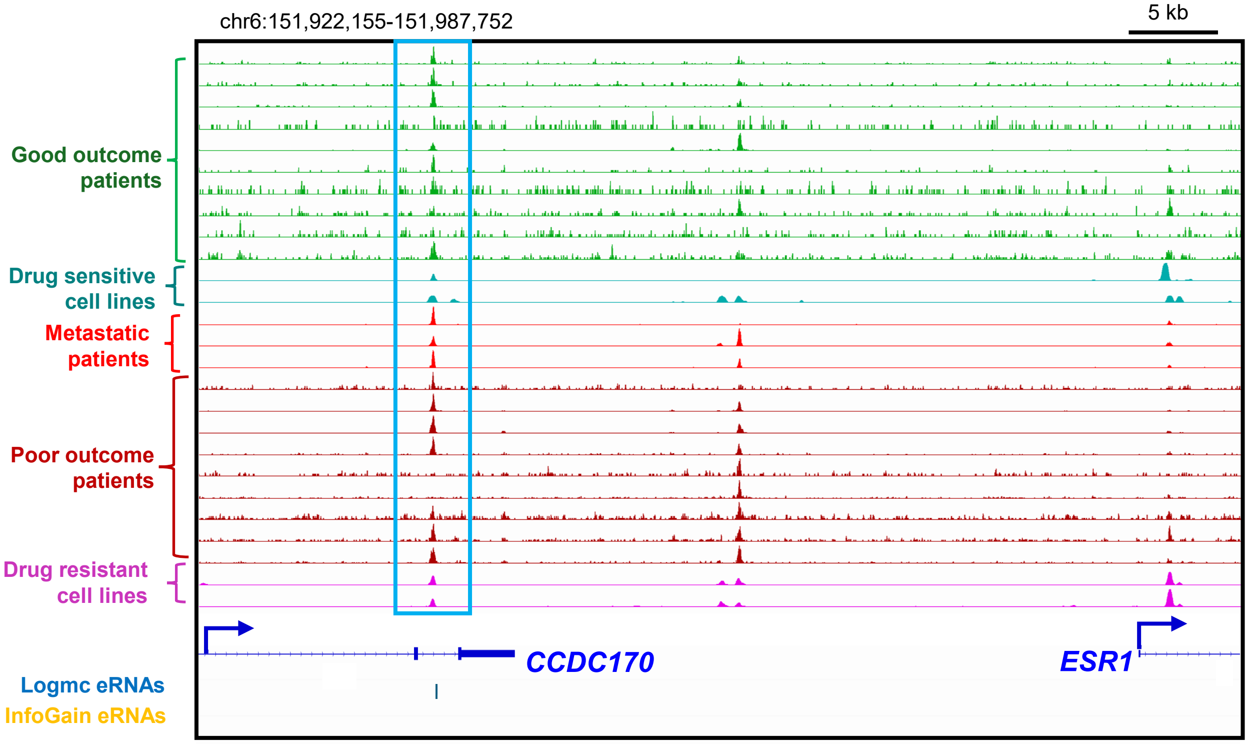


**B**

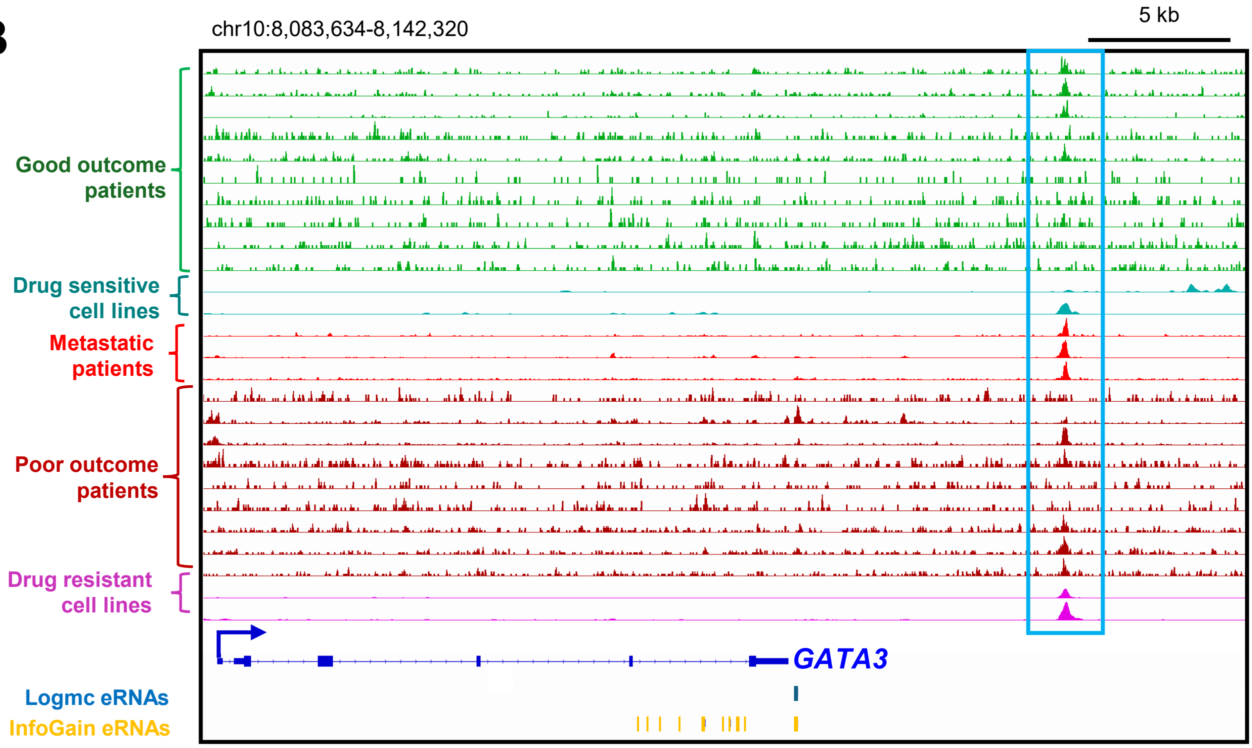

### Figure S3 cont.

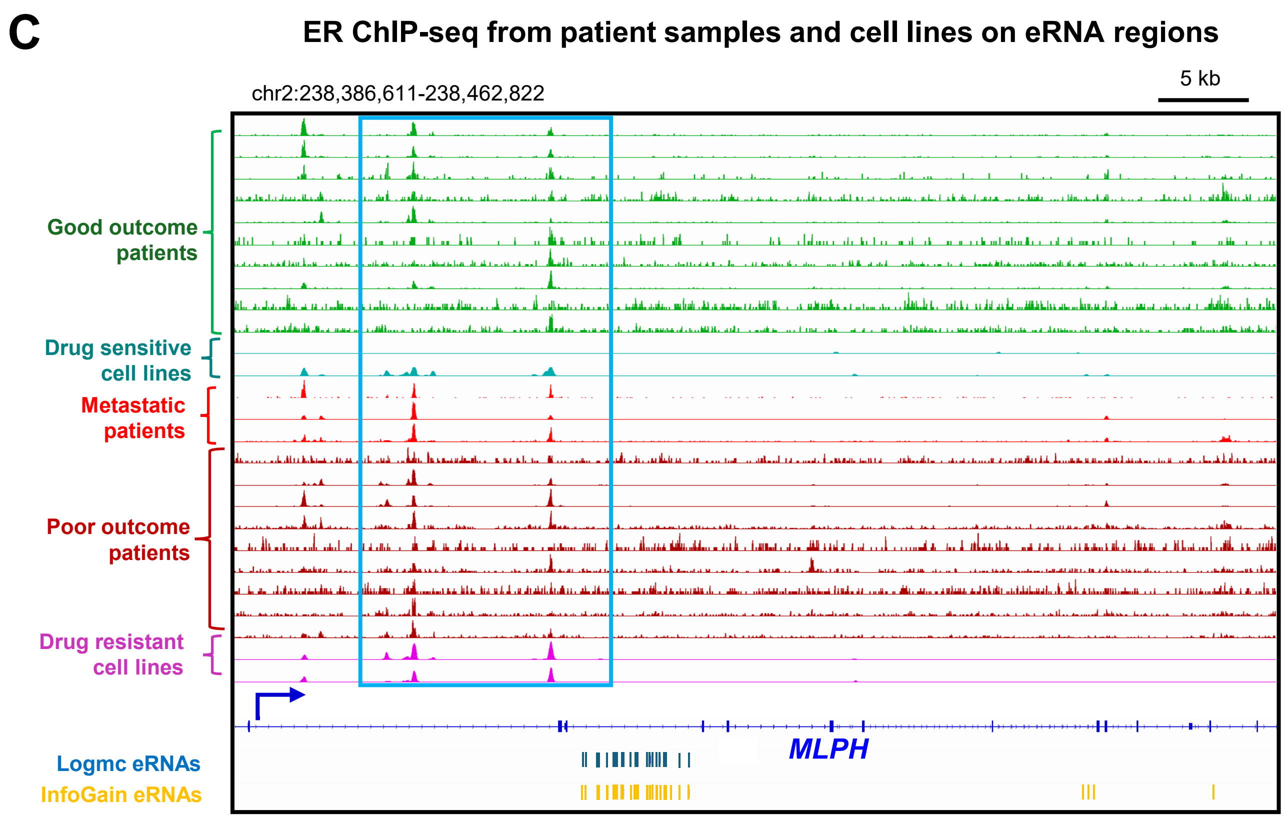


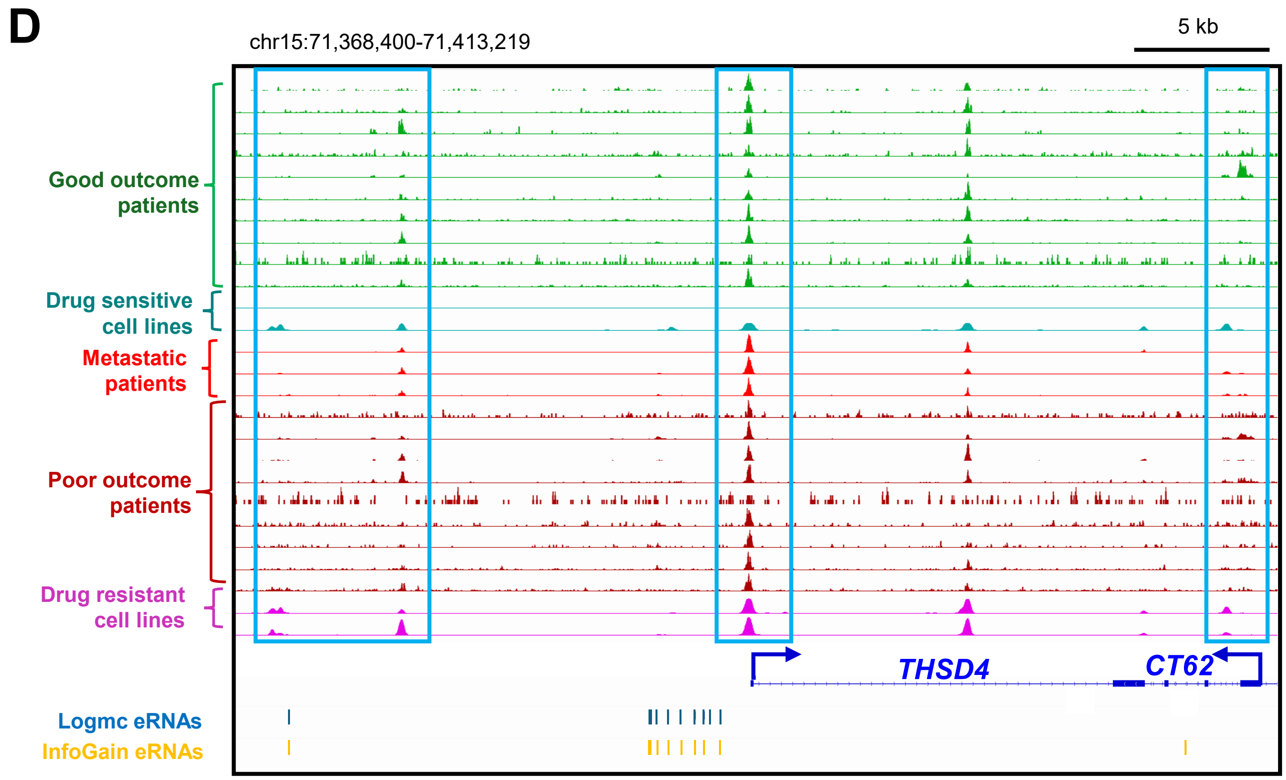

### Figure S4

**A**

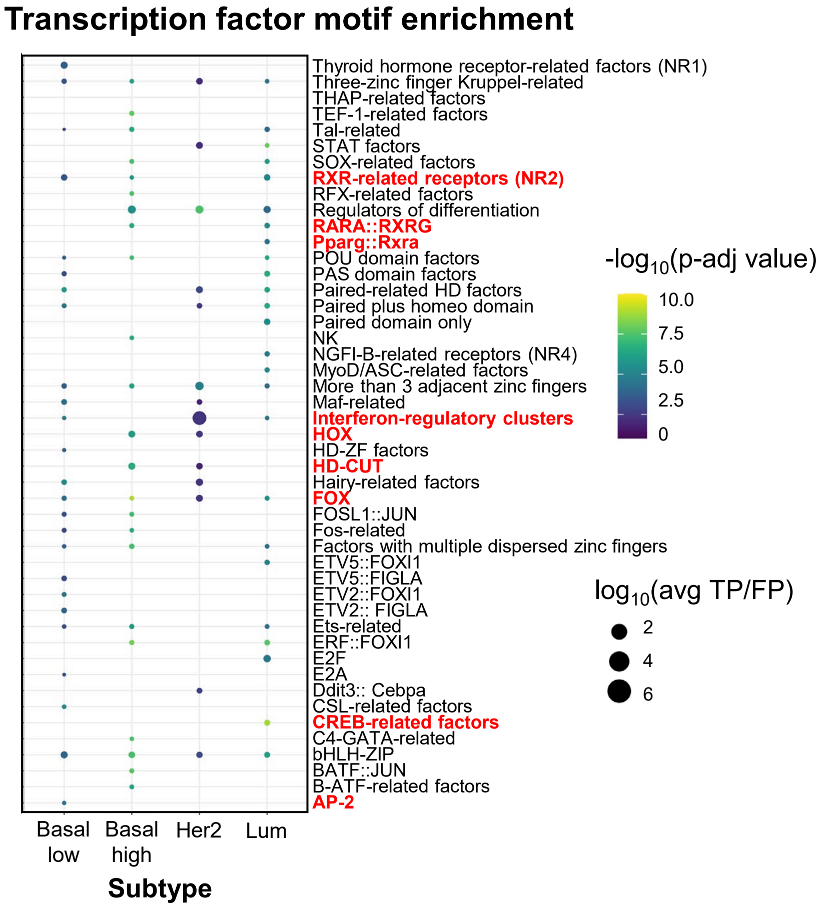
Transcription factor motif enrichment
Thyroid hormone receptor-related factors (NR1)
Three-zinc finger Kruppel-related
THAP-related factors
TEF-1-related factors
Tal-related
STAT factors
SOX-related factors
RXR-related receptors (NR2)
RFX-related factors
Regulators of differentiation
RARA::RXRG
Pparg::Rxra
POU domain factors
PAS domain factors
Paired-related HD factors
Paired plus homeo domain
Paired domain only
NK
NGFI-B-related receptors (NR4)
MyoD/ASC-related factors
More than 3 adjacent zinc fingers
Maf-related
Interferon-regulatory clusters
HOX
HD-ZF factors
HD-CUT
Hairy-related factors
FOX
FOSL1::JUN
Fos-related
Factors with multiple dispersed zinc fingers
ETV5::FOXI1
ETV5::FIGLA
ETV2::FOXI1
ETV2:: FIGLA
Ets-related
ERF::FOXI1
E2F
E2A
Ddit3:: Cebpa
CSL-related factors
CREB-related factors
C4-GATA-related
bHLH-ZIP
BATF::JUN
B-ATF-related factors
AP-2
Basal low
Basal high
Her2
Lum
Subtype
-log10(p-adj value)
10.0
7.5
5.0
2.5
0
log10(avg TP/FP)
2
4
6

**B**

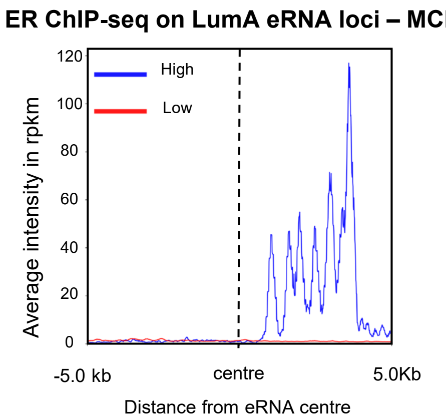
ER ChIP-seq on LumA eRNA loci – MCF7
High
Low
120
100
80
60
40
20
0
Average intensity in rpkm
-5.0 kb
centre
5.0Kb
Distance from eRNA centre

### Figure S5

**A** InfoGain-derived Her2 survival-specific eRNAs

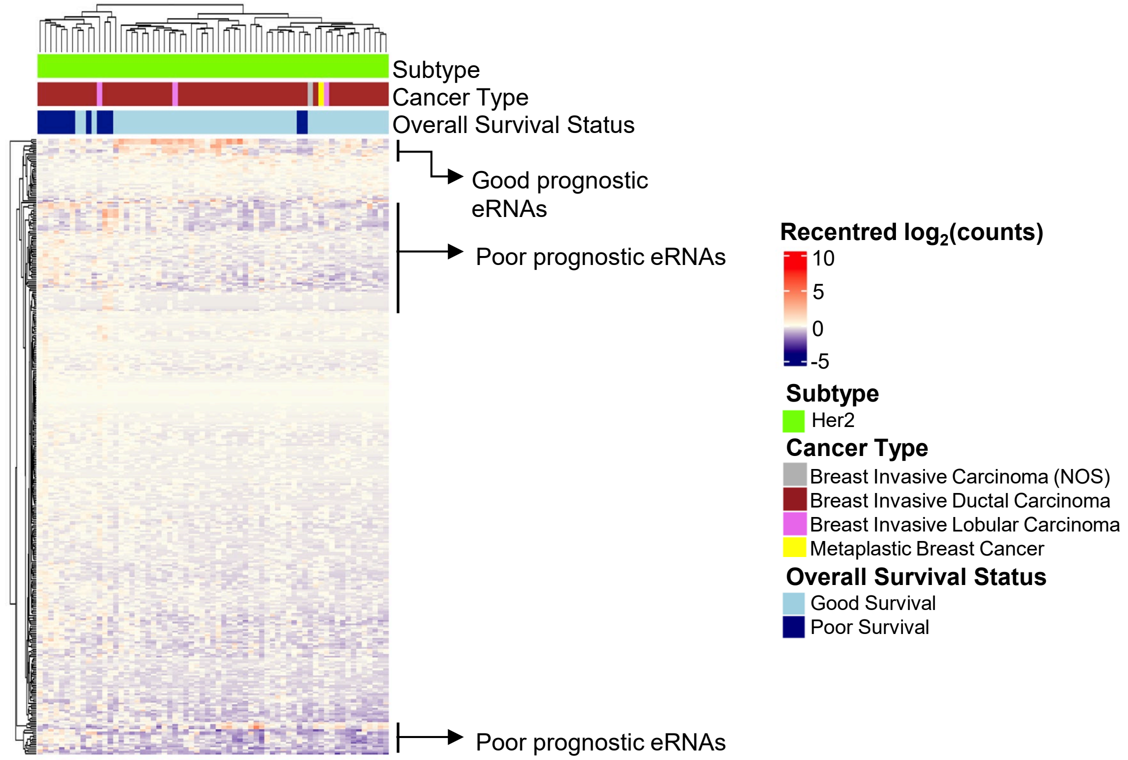


**B**

| Metrics | Good outcome | Poor outcome |
|---|---|---|
| **Precision** | 0.941 | 0.833 |
| **Sensitivity** | 0.941 | 0.833 |
| **Specificity** | 0.857 | 0.944 |
| **Accuracy** | 0.957 | 0.957 |
| **F-measure** | 0.941 | 0.833 |

**C** Prognostic eRNAs- all subtypes (Chen et al., 2018)

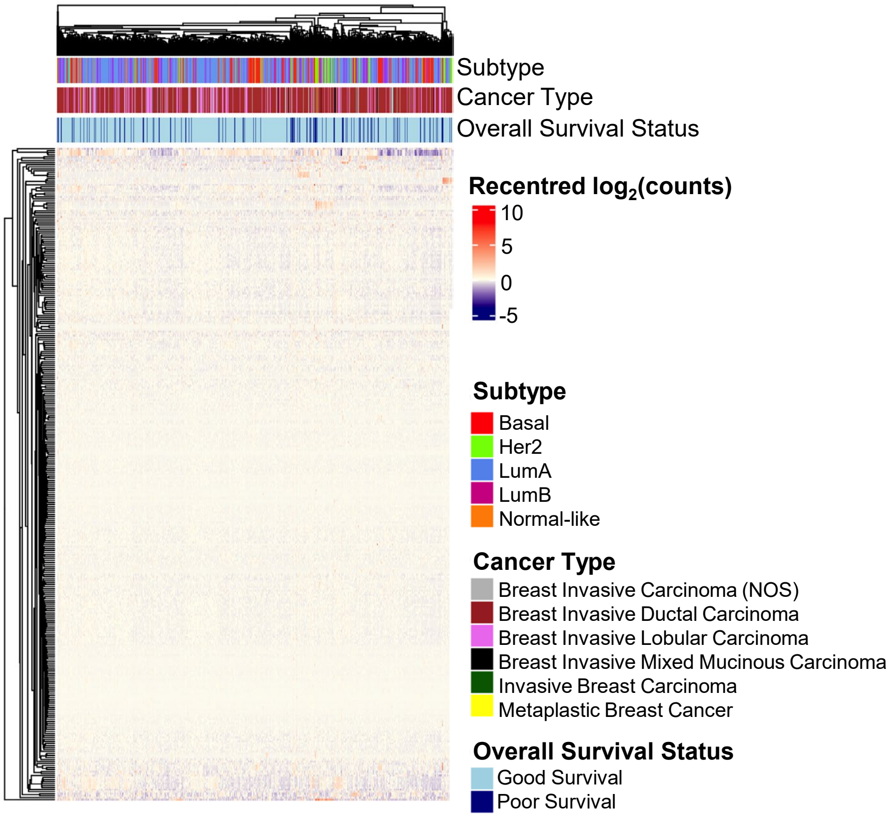
